# Synergistic effects of proteinaceous pheromone and nitrogen starvation on male gametogenesis in the anisogamous volvocine alga *Eudorina*

**DOI:** 10.1101/2025.05.26.656211

**Authors:** Hiroko Kawai-Toyooka, Makoto Ono, Takashi Hamaji, Hisayoshi Nozaki, Masafumi Hirono

## Abstract

In volvocine algae, gametogenesis is triggered by different cues depending on the species and their sexual systems. In isogamous unicellular organisms such as *Chlamydomonas reinhardtii*, which produce gametes of equal size and morphology, nitrogen depletion induces gametogenesis. In contrast, in oogamous multicellular species of the genus *Volvox*, which produce large, immobile eggs and small motile sperm, male gametogenesis is induced by a sex pheromone secreted by sperm packets (SPs), i.e., bundles of male gametes. *Eudorina*, an anisogamous multicellular volvocine alga that produces motile gametes of different sizes, is known to form SPs under nitrogen-depleted conditions. Intriguingly, a pheromone-like factor, present in male conditioned medium (CM), has also been implicated in SP formation. To clarify the relative contributions of nitrogen starvation and pheromone signaling, we conducted semi-quantitative analyses of SP induction using synchronously cultured male colonies of *Eudorina*. When CM was added to male cultures during an early maturation stage, most colonies formed SPs regardless of nitrogen availability. However, when the CM was diluted 100- to 100,000-fold, SP formation was significantly more efficient under nitrogen-depleted conditions than under nitrogen-replete conditions. Notably, SPs never formed without the addition of CM, even in a nitrogen-depleted medium. The SP-inducing activity of the CM was found to markedly decrease with protease treatment. These findings suggest that spermatogenesis in *Eudorina* is induced by a proteinaceous sex pheromone secreted by male colonies, and that nitrogen depletion, while not essential, enhances this pheromone activity.

## Introduction

Mating of complementary haploid cells (gametes) is a fundamental step in sexual reproduction across all major eukaryotic lineages [1]. Based on gamete morphology, mating systems are generally classified into three types: isogamy, where the two gametes are the same size; anisogamy, where smaller male gametes fuse with larger female gametes; and oogamy, where small, motile sperm fertilize large immotile eggs. Isogamy is considered the ancestral form, while anisogamy likely evolved, eventually giving rise to oogamy through increasing differentiation between male and female gametes [2].

The volvocine lineage serves as an excellent model system for studying the evolution of gamete dimorphism, as its members represent a range of reproductive strategies within a relatively recent divergence (Fig 1) [3–5]. For example, *Chlamydomonas reinhardtii,* a unicellular isogamous species, produces *plus* and *minus* mating-type gametes in response to nitrogen starvation [6]. Similarly, *Gonium pectorale*, an isogamous colonial species with 8 or 16 cells, undergoes gametogenesis under nitrogen-deficient conditions [7] or after prolonged cultivation [8, 9]. Thus, apparently nitrogen starvation is a key environmental cue for gametogenesis in isogamous volvocine species. In contrast, both male and female gametogenesis in the oogamous genus *Volvox* relies on sex pheromones secreted by preexisting sperm packets (SP), bundles of male gametes [10, 11]. The pheromones are highly potent; for example, a pheromone in *V. carteri*, a ∼32-kDa glycoprotein, can induce gametogenesis at concentrations as low as <10^-16^ M [10, 12–14]. These examples suggest that, during the evolutionary transition from isogamy to oogamy in the volvocine lineage, the mechanism of gametogenesis induction also shifted from a system based on environmental nitrogen resources to one relying on specific pheromones. This raises the intriguing possibility that the evolution of sexual dimorphism in volvocine green algae is linked to the change in the induction mechanism of gametogenesis.

**Fig 1.**
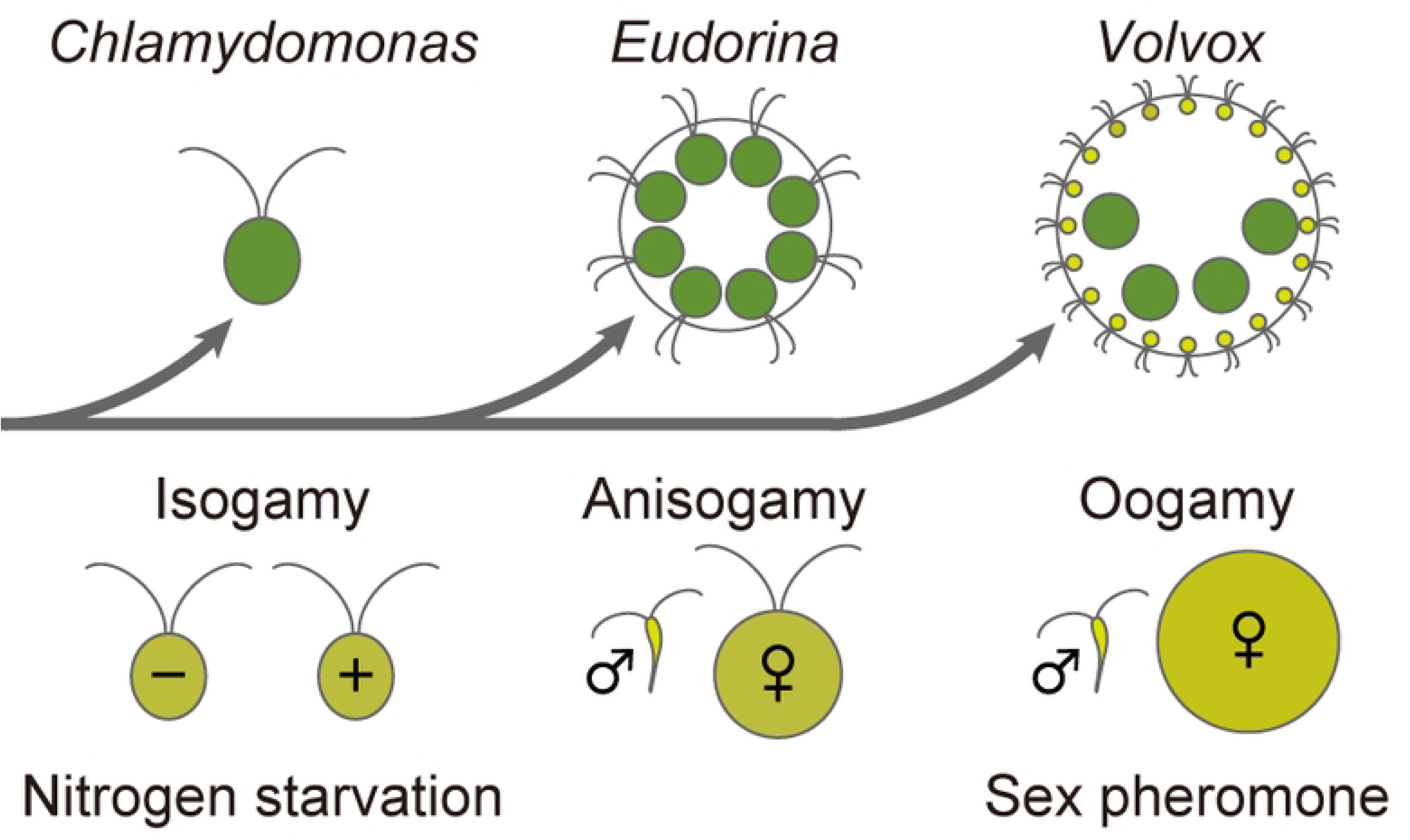
Evolutionary transitions of gamete dimorphism and gametogenesis induction cues in volvocine algae. Gamete types are illustrated for three representative volvocine genera: *Chlamydomonas* exhibits isogamy with *minus* (−) and *plus* (+) mating types, *Eudorina* shows anisogamy with motile male (♂; sperm) and female (♀) gametes, and *Volvox* displays oogamy with motile male (♂; sperm) and immotile female (♀; egg) gametes. The phylogenetic relationships among these genera are based on [15]. In *Chlamydomonas*, gametogenesis is induced by nitrogen starvation, whereas in *Volvox*, it is induced by a sex pheromone [6, 10].

*Eudorina*, an anisogamous colonial genus in the volvocine lineage, offers a valuable system for investigating the relationship between the gamete dimorphism and the gametogenesis regulation. Colonies of male and female genotypes—each comprising 16 or 32 cells—are morphologically indistinguishable during vegetative life cycle. However, upon entering the sexual reproduction process, male colonies differentiate into SPs with distinct morphology. This SP formation can be induced by culturing the colonies under nitrogen-starved conditions [16, 17]. Nevertheless, a previous study on *Eudorina elegans* reported that the addition of conditioned medium (CM) from a dense male culture also accelerated SP formation in a fresh male culture [18]. Therefore, the relative contributions of nitrogen starvation and sex pheromone to gametogenesis in *Eudorina* remain unclear.

To better understand how evolutionary changes in gamete morphology relate to the shift in gametogenesis regulation within the volvocine lineage, it is essential to clarify the relative contributions of “*Chlamydomonas*-type” nitrogen starvation and “*Volvox*-type” pheromone signaling in male gametogenesis of *Eudorina*. In this study, we investigated the effects of the male-derived CM and nitrogen starvation on male gametogenesis in *Eudorina*. Our results suggest that, at the anisogamous stage in the volvocine lineage, gametogenesis is predominantly induced by pheromone signaling, with nitrogen starvation playing a limited supporting role.

## Results

### Synchronous culture of *Eudorina*

To investigate the effects of the CM and nitrogen starvation on gametogenesis in *Eudorina*, we established cultures with a synchronized life cycle, as SP formation is closely linked to developmental stage (Fig 2A, [18]). Although a previous study showed that synchronous culture of *Eudorina* can be achieved under a 16-h light/8-h dark condition at low population density [19], light-dark cycle alone was insufficient to synchronize the colony development for the strain used in this study [20]. To overcome this, we manually isolated colonies at the pre-hatching stage—a morphologically distinct stage easily identifiable under a microscope (Fig 2B). When these colonies were incubated in a medium containing both carbon and nitrogen sources under a 16-h light/8-h dark cycle, most daughter colonies hatched from the mother colonies within 1 h of culture. The hatched colonies grew synchronously (Fig 2CD), reaching again the pre-hatching stage 25 h after colony isolation (Fig 2E). Approximately 80-98% of the colonies released next-generation colonies within 30 h. These results demonstrate that our culture method achieves sufficient synchronization of colony development for investigation of SP-inducing factors.

**Fig 2.**
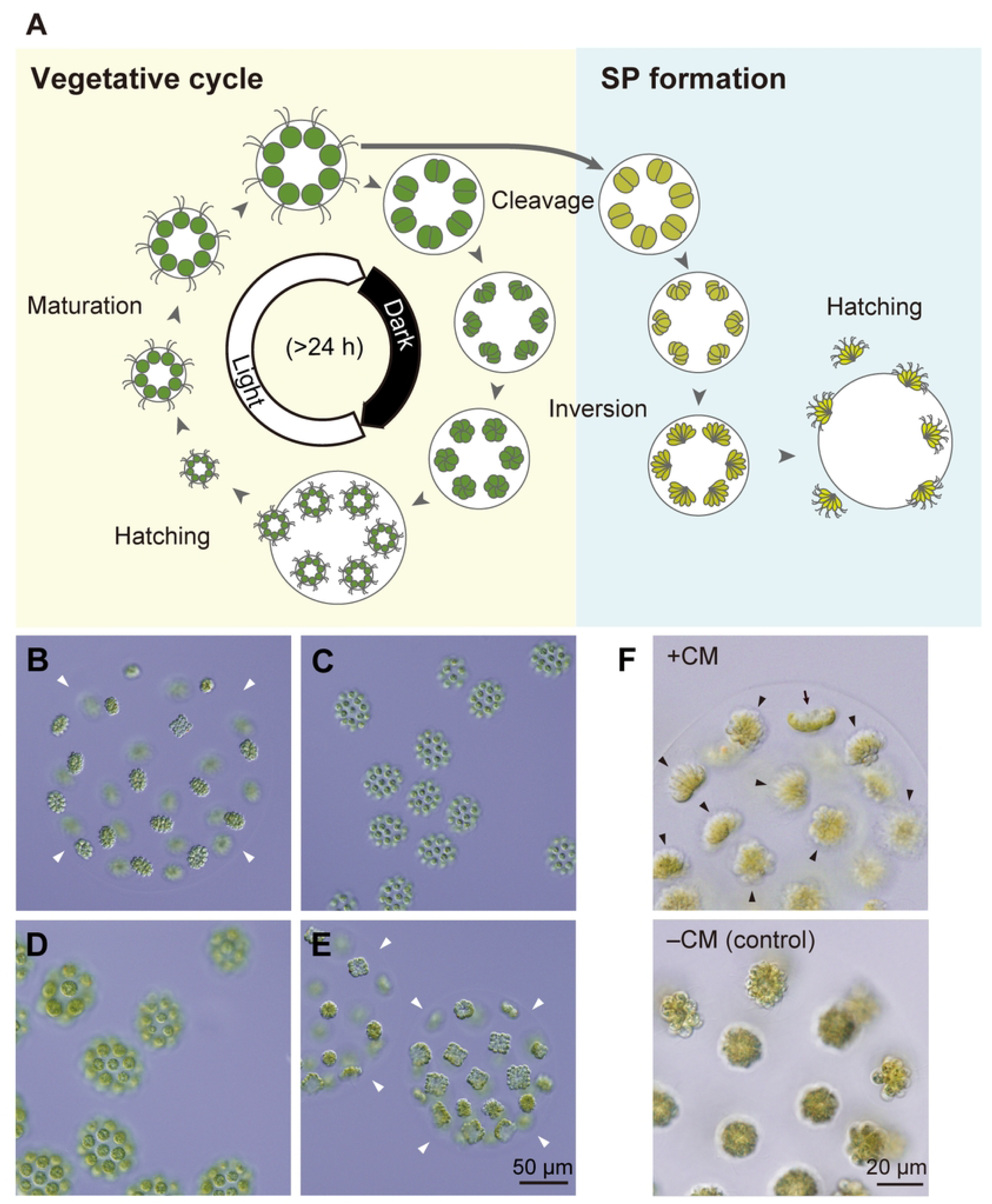
**Synchronization of the vegetative cycle and SP formation in *Eudorina***. **(A)** Schematic representation of the vegetative cycle (yellow-shaded area) and SP formation (blue-shaded area). Newly hatched colonies from the parental colony grow without cell division during the maturation stage. During the cleavage stage, cells within the colony undergo four or five rounds of division. The resulting cell clusters (embryos) then undergo inversion to form spherical embryos. These embryos further develop and reach the prehatching stage. Under a 16-h light/8-h dark cycle, the pre-hatching stage coincides with the beginning of the light period, while the cleavage stage approximately corresponds to the start of the dark period. In the SP formation pathway, embryos undergo inversion and subsequently develop into SPs. **(B-E)** Vegetative colonies (VCs) of the *Eudorina* male strain under synchronous growth conditions. **(B)** A pre-hatching colony observed 1.5 h after the start of the light period. Colonies at this stage were isolated for synchronous culture. **(C, D)** Colonies at the maturation stage observed at 6 h **(C)** and 17 h **(D)** after the start of the light period, with all colonies developing synchronously. **(E)** Pre-hatching colonies reappeared at 26.5 h. Arrowheads indicate the edges of the parental colonies’ extracellular matrix. **(F)** SP formation induced by the addition of male CM. Male colonies grown under synchronous conditions were incubated with male-derived CM for approximately 24 h. Bouquet-shaped SPs (arrowheads) were observed within the parental colonies, along with a pre-inversion embryo (arrow) that could not yet be identified as either VC or SP. As a control, when CM was not added (–CM), only spherical VCs were observed. The male strain used in **(B-F)** was 2017-0525-EF2-15.

### Proteinaceous factor(s) in male CM induce SP formation

Using synchronized cultures, we evaluated the SP-inducing activity of the CM obtained from a high-density culture of male colonies grown asynchronously for 3-5 days [21]. The CM was added to a synchronized male culture at the start of the light period (corresponding to the pre-hatching stage or a stage slightly after hatching), making up one-third of the total medium volume. Approximately 60% of the colonies differentiated into bouquet-shaped SPs within ∼24 h of the CM addition, whereas no SP was formed in the control culture without CM (Fig 2F). These results clearly demonstrate that the male-derived CM contains factor(s) that induce SP formation, and that nitrogen starvation is not required for the induction when sufficient CM is added.

In *V. carteri*, gametogenesis is induced by a proteinaceous sex pheromone present in the male CM [13]. To determine whether the SP-inducing factor(s) in the male CM of *Eudorina* is similarly proteinaceous, we tested the activity of CM after treating it with various concentrations of pronase, a protease mixture known to degrade the *Volvox* pheromone [22], with or without heat treatment (Fig 3). Heat treatment alone reduced SP formation from ∼87% to ∼30%. Pronase treatment further decreased activity in a dose-dependent manner (Fig 3). These results suggest that, as in *V. carteri*, the SP-inducing factor(s) in the male *Eudorina* CM is also proteinaceous.

**Fig 3.**
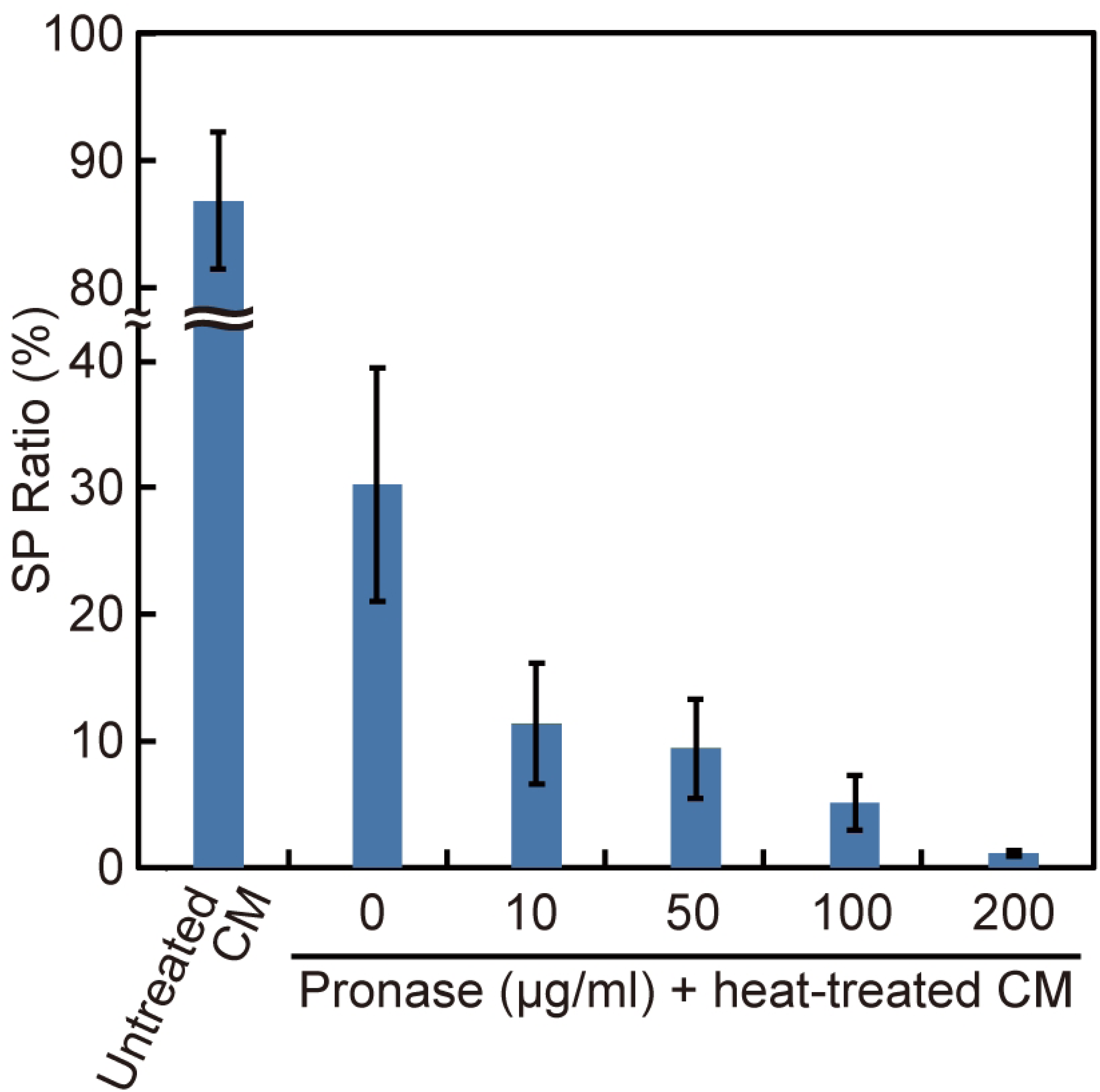
**Effect of pronase pretreatment of male CM on SP formation**. Male CM was treated with pronase of the concentrations indicated on the abscissa, heat-inactivated (80°C, 20 min), and then applied to male colonies at a final volume of 10%. As a control, untreated CM (without pronase and heat treatment) was applied at the same volume. The SP ratio (%) was calculated as the percentage of SPs relative to the total number of colonies (SPs and VCs) in each sample. Each value represents the mean ± standard deviation (SD) from three biological replicates (≥90 colonies in total). The experiment was independently repeated, yielding consistent results. The male strain used was 2022-1122-EF4-M1.

### Sensitivity to CM depends on the developmental stages

For SP-formation in *V. carteri* to occur, its reproductive cells, gonidia, must be exposed to the sex pheromone 6–8 hours prior to the start of cell division [14]. To investigate whether *Eudorina* exhibits a similar sensitive period in its life cycle, we next examined the responsiveness of synchronously cultured male colonies to the added CM at various developmental stages. Following hatching, the colonies underwent a period without cell division during the light phase. We call this stage maturation stage after the corresponding stage in the *Volvox* life cycle ([10]; S1 Fig). This stage was followed by cell division during the dark phase (the cleavage stage; Fig 2A). After cleavage, the resulting cell clusters within the colony underwent inversion to assume inside-out spherical forms, which would mature into the next-generation colonies. When colonies from the prehatching stage up to the middle of the maturation stage (∼9 h after cultivation began) were treated with the CM, SP formation rate was ∼90% (Fig 4). However, when the CM treatment was performed on colonies in the latter half of the maturation stage (∼12 h after cultivation began), the SP formation rate decreased to ∼60%. Once the colonies had entered the cell cleavage stage, no SP formation was observed. These results indicate that *Eudorina* cells must be exposed to the SP-inducing factors in CM at least 6-8 h before the start of cell division to differentiate into SP.

**Fig 4.**
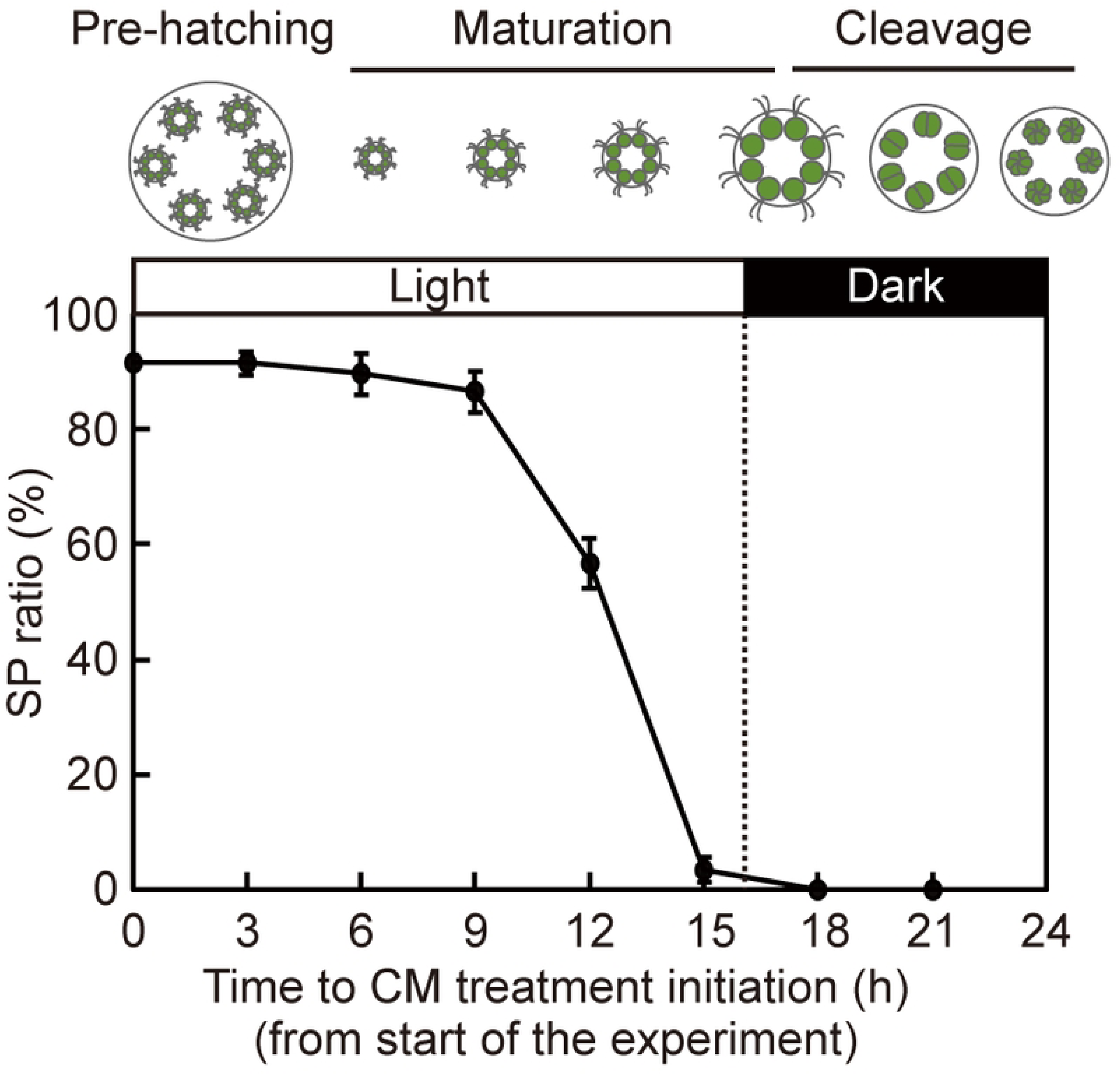
**Time course of CM-induced SP formation efficiency during colony development of *Eudorina***. Male colonies at various life cycle stages were treated with CM at a final volume of 10%, and the SP ratios (%) were quantified. Each value represents the mean ± SD from three biological replicates (≥68 colonies in total). Schematic illustrations above the graph depict the developmental stages of the colonies at each time point, along with the light conditions (16-h light/8-h dark). The male strain used was 2022-1122-EF4-M1.

### Nitrogen starvation promotes SP formation in diluted CM

We next examined the influence of nitrogen starvation on the SP formation efficiency after the addition of diluted CM. When 1/10 volume of CM had been added to the culture, the SP formation efficiency did not noticeably change depending on nitrogen availability (Fig 5). However, as the amount of CM was further reduced, the SP formation efficiency tended to increase in the nitrogen-deficient medium. Promotion by nitrogen deficiency became more pronounced at lower CM concentrations. With the CM volume reduced to 1/100,000 of the culture volume, SP formation occurred only in the nitrogen-deficient medium. SP formation did not occur in the complete absence of CM (Fig 5). These results indicate that while nitrogen starvation alone cannot induce SP formation, it significantly enhances CM activity when CM concentration is low.

**Fig 5.**
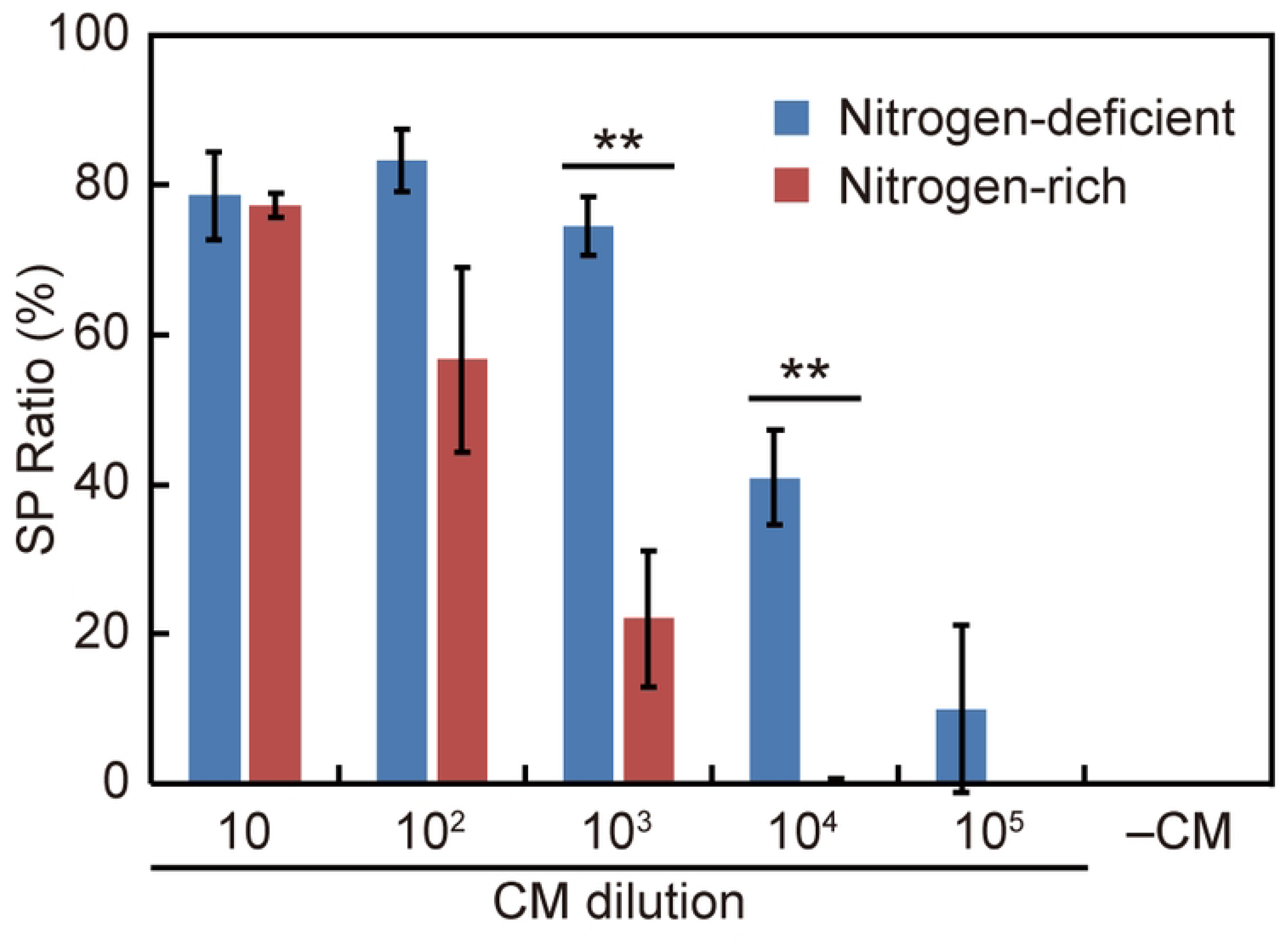
**Effect of CM dilution and nitrogen availability on male CM-induced SP formation** SP ratio (%) was measured after applying serially diluted CM (10–10⁵) to male colonies in either nitrogen-deficient (blue) or nitrogen-rich (red) medium. Each value represents the mean ± SD from three biological replicates (≥68 colonies in total). Asterisks indicate p < 0.01 based on an independent two-sample, two-tailed t-test. The experiments were independently repeated and essentially the same results were obtained. The male strain used was 2022-1122-EF4-M1.

## Discussion

This study demonstrates that in *Eudorina*, an anisogamous volvocine alga, a proteinaceous sex pheromone present in the male CM is the primary inducer of SP formation, and that this effect is enhanced by nitrogen starvation. Although the SP-inducing activity in the CM has already been reported in *Eudorina elegans* [18], our current study has, for the first time, shown that the responsible factor is proteinaceous. These findings are particularly intriguing given *Eudorina*’s intermediate evolutionary position within the volvocine lineage—between *Chlamydomonas,* where nitrogen starvation is the primary inducer, and *Volvox*, where sex pheromones play the major role (Fig 1) [6, 10, 15, 17, 23, 24].

The cooperative effect of sex pheromone and nitrogen starvation on SP formation may also be conserved in *Pleodorina*, another anisogamous member of the volvocine lineage. This possibility is supported by previous studies showing that SP formation in *Pleodorina* (*P. japonica* and *P. starrii*) was induced by culturing the colonies to high density, followed by a reduction in the nitrogen concentration [25–28]. These conditions for SP formation may reflect synergistic effects of sex pheromone and nitrogen depletion, as we demonstrated with *Eudorina* in the present study, suggesting that the mechanism underlying SP formation is shared among various species of anisogamous volvocine algae.

In *Volvox*, the effect of nitrogen starvation on SP formation has not been reported. In *V. carteri*, SP formation is induced by treating vegetative colonies (VCs) with male CM containing the sex pheromone, even in the presence of sufficient nitrogen sources [29–31]. In other *Volvox* species, such as *V. ferrisii*, *V. africanus,* and *V. reticuliferus*, gametogenesis is spontaneously induced during continuous cultivation in nitrogen-containing medium [32–36]. Thus, gametogenesis in all studied *Volvox* species tends to occur under relatively nitrogen-rich conditions, suggesting that nitrogen starvation is unlikely to promote sexual induction in oogamous volvocine algae. In contrast, in *Chlamydomonas*, sex pheromones have not been reported to play a role in gametogenesis [6, 37].

Taken together, these previous findings and our results here allow us to classify the gamete inducers in volvocine algae into three types: the “*Chlamydomonas*-type,” in which nitrogen starvation alone serves as the trigger; the “*Eudorina*-type,” in which both sex pheromones and nitrogen starvation act cooperatively; and the “*Volvox*-type,” in which sex pheromones alone induce gametogenesis. Since these three types correspond to the three modes of sexual reproduction modes (isogamy, anisogamy, and oogamy), we can imagine that the mechanism of gametogenesis induction and the gamete morphology may have evolved in close association.

Our preliminary observation showed that the sex pheromone of *Volvox* is unable to induce SP formation in *Eudorina* (no SPs were observed in over 200 colonies counted, n=3). Therefore, pheromone signaling in these organisms appears to be species-specific. However, there are some similarities in the pheromone actions in the two species. In *Volvox*, cells within a colony are differentiated into two types, somatic and reproductive. When the sex pheromone acts on gonidia, they undergo differentiation into SPs through an intermediate state called androgonidia ([10]; S1 Fig). For this differentiation to occur, gonidia must be exposed to the sex pheromone 6–8 hours before the start of cell division [14]. This temporal requirement is similar to the pheromone sensitivity period in *Eudorina*, where vegetative cells must be exposed to the sex pheromone 6–8 hours prior to the initiation of cell division (Fig 4). Thus, despite the difference in the type of cells that differentiate into SPs, i.e., vegetative cells in *Eudorina* and gonidia in *Volvox,* the timing of pheromone sensitivity appears similar between these organisms.

In summary, our study demonstrates the synergetic effects of sex pheromones and nitrogen starvation on SP formation in *Eudorina*, thereby positioning this species as a significant evolutionary link between *Chlamydomonas* and *Volvox*. The categorization of gamete inducers into three types—*Chlamydomonas*-type, *Eudorina*-type, and *Volvox*-type—suggests a correlation between gametogenesis induction mechanisms and sexual reproduction modes. These findings provide valuable insights into the evolutionary transition of sexual induction mechanisms in colonial algae.

## Materials and methods

### Strains

Strains used were progenies of the *Eudorina* strains 2006-703-Eu-15 (male, NIES-2735) and 2006-703-Eu-14 (female, NIES-2734), originally collected in Taiwan [17]. Because long-term subculturing led to a decline in SP formation efficiency in the male strain, we maintained male strains with high SP formation efficiency through repeated crossing of existing strains. The strains used for analyses were as follows: 2010-623-F1-E3 (NIES-4100; [21]), an F1 male strain resulting from the initial parental cross; 2017-0525-EF2-15 (NIES-4564) and 2017-0525-EF2-21 (NIES-4565), male and female strains, respectively, generated from a cross between the F1 male strain and the female parental strain; 2021-0602-EF3-M6 (NIES-4567) and 2021-0602-EF3-M4 (NIES-4568), male and female strains, respectively, generated from a cross between 2017-0525-EF2-15 and 0525-EF2-21; 2022-1122-EF4-M1, a male strain generated from a cross between 2021-0602-EF3-M6 and 2021-0602-EF3-M4.

### Culture

*Eudorina* was maintained in 18 x 150 mm screw-cap tubes containing 10 mL of artificial freshwater-6 (AF-6) medium [38], with ∼300 µL of culture transferred to fresh medium every four weeks. Cultures were grown at 25 ± 2°C under a 14-h light/10-h dark or 12-h light/12-h dark cycle, illuminated by cool white fluorescent and/or LED lamps at an intensity of <100 μmol photons m^−2^ s^−1^.

For synchronous growth of VCs, ∼30 µL of stock culture was transferred to a 60-mm Petri dish containing 10 mL of nitrogen-rich medium (VTAC; [38]; S1 Table). and incubated for 3 days at 32 ± 1.5°C under a 16-h light/8-h dark cycle, with illumination from cool white fluorescent and/or LED lamps at 230–260 μmol photons m⁻² s⁻¹. At the beginning of the light period, 10 pre-hatching colonies were selected under a binocular dissecting microscope, washed with VTAC medium to remove any residual CM, and then cultured in 3 mL of fresh VTAC medium in a 30-mm Petri dish for the indicated time.

### CM preparation

For small-scale CM preparation, ∼400 µL of stock culture of the male strain maintained for ∼4 weeks was transferred to 10 mL of VTAC in a 60-mm Petri dish and cultured at 25 ± 2°C under a 14-h light/10-h dark cycle with a light intensity of 180–280 μmol photons m⁻² s⁻¹ for 3 days. The culture was then diluted with twice the volume of nitrogen-deficient medium (mating medium; [25]; S1 Table) and further incubated for an additional 1–2 days.

For large-scale CM preparation, ∼40 mL of the stock culture of the male strain was transferred to 300 mL of VTAC in a 500 mL flask. The culture was incubated under aeration at 32 ± 1.5°C under a 16-h light/8-h dark cycle with a light intensity of 170–220 μmol photons m⁻² s⁻¹ for 3 days.

After confirming SP formation, the culture was centrifuged, and the supernatant was filtered through a 0.22-µm filter to remove cells and debris. The resulting CM was stored at -80°C. For the experiment shown in Fig 5, 1 mL of large-scale prepared CM was transferred into a 14 kDa-cut-off dialysis membrane (Viskase) and dialyzed against 1.8 L of 10 mM HEPES buffer (pH 7.8) overnight at 4°C before use.

### SP formation assay

Pre-hatching VCs were selected and washed with either VTAC or mating medium. A total of 8–20 colonies were then transferred to each well of a 12-well plate containing 2 mL of the respective medium. Alternatively, ∼26 µL of a suspension containing ∼100 freshly hatched colonies, prepared under synchronous growth conditions, was added to each well. Each well was supplemented with CM or with either distilled water or 10 mM HEPES buffer (pH 7.8) as a negative control. For the experiment shown in Fig 4, 8 pre-hatching colonies were transferred to each well simultaneously, and CM was added to different wells at 3-h intervals. The plates were incubated at 32°C under a light intensity of 280–380 µmol photons m⁻² s⁻¹ with a 16-h light/8-h dark cycle for 24–30 h, until most SPs and VCs had hatched from the mother colonies. Cultures were then fixed by adding glutaraldehyde to a final concentration of 0.25% (v/v). SPs and VCs were counted in 200 µL of the culture medium. The SP ratio was calculated as the proportion of SPs relative to the total number of colonies. All experiments were independently repeated, and representative results are presented.

### Pronase treatment

Small-scale CM prepared from the male strain was incubated with the indicated concentration of pronase (#10165921001; Roche) at 37°C for 1 h. To inactivate the pronase, the mixture was heated to 80°C for 20 min. After cooling, 200 µl of the mixture was subjected to the SP formation assay with mating medium.

### Light microscopy

Microscopic images were captured using an upright microscope (BX-53; Olympus) equipped with UPlanSApo 40X (NA 0.95) or 100X (NA 1.40) objective lenses (Olympus) and differential interference contrast (DIC) optics. Images were recorded with a DP71 camera (Olympus) controlled by DP controller software (version 1.2.1108; Olympus) and processed with Photoshop CC (version 20.0.8; Adobe).

## Acknowledgments

We thank Ritsu Kamiya for critically reading the manuscript. We also thank Yumi Asano for technical assistance, Kohei Takahashi for sharing mating techniques, and Yoshiki Nishimura for experimental support. We are grateful to Takako Kato-Minoura for her continued encouragement and support.

## Supporting Information

**S1 Figure. Schematic representation of the vegetative cycle and sperm packet (SP) formation in the *Volvox carteri* male strain (for comparison)**. In *Volvox*, the vegetative cycle (yellow-shaded area) takes ∼48 h under a 16-h light/8-h dark cycle, compared to ∼24 h in Eudorina. This longer duration is due to the prolonged expansion phase, during which both the parental colony and its enclosed daughter colonies expand for ∼24 h after inversion. When gonidia sense the sex pheromone 6–8 h before the cleavage stage, they undergo modified type of embryogenesis to form a colony containing androgonidia (blue-shaded area). Subsequently, the androgonidia divide and form SPs through incomplete inversion.

**S1 Table. Composition of VTAC and mating medium based on [S1, S2].**

